# A non-local model for cancer stem cells and the tumor growth paradox

**DOI:** 10.1101/019604

**Authors:** I. Borsi, A. Fasano, M. Primicerio, T. Hillen

## Abstract

The *tumor growth paradox* refers to the observation that incomplete treatment of cancers can enhance their growth. As shown here and elsewhere, the existence of cancer stem cells (CSC) can explain this effect. CSC are less sensitive to treatments, hence any stress applied to the tumor selects for CSC, thereby increasing the fitness of the tumor. In this paper we use a mathematical model to understand the role of CSC in the progression of cancer. Our model is a rather general system of integro-differential equations for tumor growth and tumor spread. Such a model has never been analysed, and we prove results on local and global existence of solutions, their uniqueness and their boundedness. We show numerically that this model exhibits the tumor growth paradox for all parameters tested. This effect becomes more relevant for small renewal rate of the CSC.

## 1 Introduction

Tumor stem cells, or cancer stem cells (CSC) have been identified in many cancers, including carcinomas of the breast, brain, colon, prostate, pancreas, ovary, as well as in sarcomas and in leukemia [5]. Stem cells in general are pluripotent and they generate cells of various cell lineages. In the hematopoietic system, for example, the progenitor cells are often classified as multipotent progenitors who give rise to a lineage of transient, transient amplifying and differentiated cells. In cancer these differentiation stages are more diffuse and a clear distinction between these stages is often not possible. Hence, to investigate the role of stem cells, we combine all non stem cancer cells into one group called cancer cells (CC). While it is not always easy to identify cancer stem cells, it is widely accepted that they are instrumental in tumor progression and control. In fact, the control or eradication of CSC has become a focus for treatment design [4].

In this article we develop and analyse a mathematical model for CSC and CC which consists of a non-linear coupled system of integro-differential equations (iDEs). Existence results for this coupled system are not readily available, hence we prove mathematical properties on positivity, boundedness, existence and uniqueness. We show numerically that this iDE model supports a *tumor growth paradox*, i.e. the fact that a tumor with larger death rate for CC might grow bigger than a tumor with lower CC death rate. We then analyse the sensitivity of the tumor growth paradox on the spatial spread rates of the tumor cells. We find that the tumor growth paradox arises for all tested parameter values. It is more pronounced in cases of low mobility of the CSC and a low renewal rate for CSC. Finally, we consider an example of incomplete radiation treatment, where the tumor after radiation grows larger then it would have grown if left untouched - the tumor growth paradox at work.

### 1.1 Modelling of cancer stem cells

Mathematical modelling is an established method to help to understand complex biological systems. Related to tumor stem cells several models have been developed recently [4,7,15,2,16,11,17]. Aspects of cancer stem cells that are of importance to cancer progression include repopulation, treatment resistance, stem cell differentiation, positive and negative feedback loops, de-differentiation mechanisms, competition for oxygen and nutrients and spatial arrangements. Mathematical modelling of all these aspects in one big model is certainly possible, but its potential use is limited. The more detail we include, the more specific assumptions and parameter values are needed. However, mathematical modelling has the advantage that sub models can be studied on their own, thereby allowing us to focus on one or two aspects at a time, which is often impossible *in vivo*. In this paper we focus on the aspect of spatial propagation and spatial crowding by cancer stem cells (CSC) and non-stem cancer cells (CC).

In this context, Enderling *et al.* [6] developed an individual based cellular automaton model, where individual cells are described by elements of a square grid. Each grid point can be either a CSC or a CC or empty. Cells are able to divide if a free grid cell is available near by, otherwise cells become dormant (quiescent). Enderling *et al.* study the effect of spatial inhibition on the simulated tumor. If the death rate for the CC is low, then CC quickly surround CSC, who in turn lose free space for replication. Hence they turn quiescent until the whole tumor stops growing (or grows very slowly). If however, the death rate for the CC compartment is increased, for example due to treatment, then CSC find open space to grow into. They produce more CSC through occasional symmetric divisions and as a result the tumor becomes bigger. The effect that increased CC death can lead to a larger tumor has been termed the *tumor growth paradox*. In fact it is commonly observed in clinical practice that incomplete treatment can lead to an increased tumor burden after treatment. Enderling *et al.* [6] point out that their result requires some movability of CSC, otherwise the tumor growth paradox would not arise.

Hillen *et al.* [11] recast the dynamics of the cellular automaton model of Enderling *et al.* [6] as a system of integro-partial differential equations (iPDE) for continuous population densities of CSC and CC. While the system was developed in Hillen *et al.* it was not analysed there. Instead, Hillen *et al.* simplified the iDE model into a (spatially homogeneous) system of ordinary differential equations (ODEs). For this ODE system they used geometric singular perturbation theory [10] to mathematically prove the existence of a tumor growth paradox. The analysis of the full iPDE model for cancer stem cells was left open and this is the topic of a recent paper by Maddalena [13] and the present paper. We discuss Maddalena’s results later in Section 1.2 in Remark (4).

In the next Section 1.2 we will motivate the iDE model for CSC and CC from biological principles. In Section 2 we will present a detailed mathematical analysis of the above model. Under rather weak assumptions we can show that continuous solutions exist, they are unique, and they depend continuously on the initial conditions. The model contains a crowding term *F* (*p*) with crowding capacity 1 and we can prove that solutions stay bounded between 0 and 1.

In Section 3 we showcase some numerical experiments of the above model. We show clearly that the tumor growth paradox arises, i.e. a tumor with a larger death rate for CC will outgrow a tumor with a lower death rate. The reason is, that a larger death rate for CC leads to a selective advantage for the CSC. Hence, a tumor with larger death rate for CC is CSC dominated, while a tumor with lower death rate is CC dominated. We study the sensitivity of the tumor growth paradox on the model parameters and we find that the tumor growth paradox is more pronounced for slow moving CSC and in cases where the CSC renewal rate is small. Larger spread of cells leads to a more homogeneous distribution and to less effect of the tumor growth paradox.

Additionally, we show some qualitative simulations for incomplete treatment and we find scenarios where the tumor after treatment will grow bigger than the corresponding untreated tumor.

This is, of course, not the final answer and the above model shows many more interesting properties in both ways, mathematically as well as for the application. In the concluding Section 4 we will discuss the relevance of our findings, and possible future avenues of investigation.

### 1.2 The iDE model for CSC and CC

To motivate our iDE (integro-differential equation) model we first recall the ODE model of Hillen *et al.* [11]

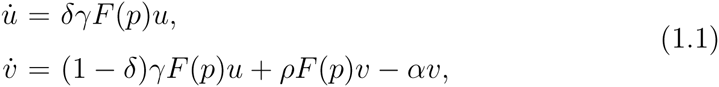

with *p* = *u* + *v*, where *u*(*t*), *v*(*t*) denote time dependent densities of CSC and CC, respectively. The parameter *γ* > 0 denotes the average mitosis rate for CSC, while *ρ* > 0 describes the mitosis rate for CC; *α >* 0 denotes the death rate of the CC cells. Death can be related to natural death or treatment induced cell death. The parameter *δ* > 0 describes the average fraction of CSC in the progeny of a CSC. There is no death term in the CSC compartment in this model, since we assume that CSC are immortal and also are less sensitive to treatment. It is an extreme case, and a small death rate for CSC could be incorporated if needed. The *F*(*p*) term describes competition for space. *F*(*p*) is a monotonically decreasing function describing the inhibitory effect of a cell density *p*. If *p* exceeds a given crowding threshold *p*\*, then *F*(*p*) = 0 for all *p* ≥ *p*\*. Here we normalize *p*\* = 1.

Hillen *et al.* proved in [11] that the above model (1.1) shows a tumor growth paradox, which, for this and more general cases we define as follows.

#### Definition 1.1

*Let p*_*α*_(*t*) *denote the total tumor size at time t >*0*, where α ≥* 0 *denotes the death rate of non-stem cancer cells (CC). The model exhibits a* **tumor growth paradox** *if there exist death rates α*_1_ < *α*_2_ *and a time interval* (*t*_1_, *t*_2_), *t*_2_ > *t*_1_ > 0, *such that p*_*α*1_ (0) = *p*_*α*2_ (0) *and*

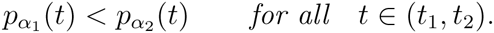

Hillen et al. [11] showed that (1.1) always has at least two steady states (0, 0) and (1, 0) and a third steady state (0, *v*_0_) if the equation *F*(*v*_0_) = *α* has a solution. The simplex *S* := {*u* ≥ 0, *v* ≥ 0, *u* + *v* ≤ 1} is positively invariant and the only global attractor in *S* is (1, 0). Hillen et al [11] used methods from geometric singular perturbation theory [10] and showed that for small *δ* > 0 there exists a slow manifold inside *S* which attracts each orbit. The slow manifolds depend monotonically on *α* in such a way that the population with the larger *α* value does grow slower hence implying the tumor growth paradox. Enderling et al [6] argued that in their cellular automaton model the tumor growth paradox needed some form of spatial random movement. Hence, in this paper here, we include spatial redistributions expressed through non-local integral terms to see if spatial movement would enhance, preserve or remove the tumour growth paradox as compared to (1.1).

To make this model spatially dependent, we use the general framework of a *birth-jump process* as introduced recently [12]. We describe spatial redistribution by an integral kernel *k*(*x, y, p*(*x, t*)). In our spatial model we assume that at mitosis one daughter cell can take the location of the mother cell, while the other daughter cell is transported to another location. This can be nearby, depending on the kernel *k*(*x, y, p*(*x, t*)). The kernel *k* is a transitional probability density in the sense that *k*(*x, y, p*(*x, t*))Δ^*n*^*x* is the probability that a daughter cell which is released at *y* settles in an n-dimensional volume element at *x* with side length Δ*x* per unit of time. This probability depends on the local density *p*(*x, t*) to describe the volume filling effect. We choose

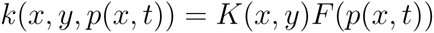

to separate redistribution *K*(*x, y*) and the crowding effect *F*(*p*). The spatial model reads:

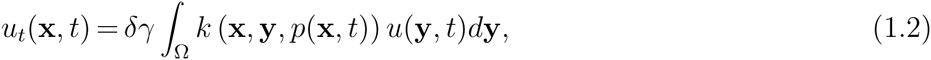

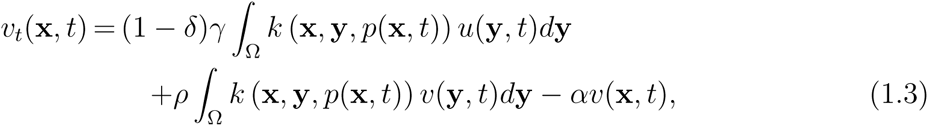

where

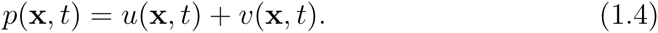

The spatial domain is denoted as Ω. The functions *u*(**x**, *t*) and *v*(**x**, *t*) represent the densities of the cancer stem cells (CSC) and of the non-stem cancer cells (CC), respectively. The parameters *γ* > 0 and *ρ* > 0 are the replication rates of the two families of cells; *δ*, 0 *<* δ < 1, is the average fraction of CSC in the progeny of a CSC; *α* is the mortality rate of the differentiated cells, and *k*(**x**, **y**, *p*(**x**, *t*)) is the probability density that a cell located at **y** generates a cell at **x**. We assume *k*(**x**, **y**, *p*(**x**, *t*)) = *K*(**x**, **y**)*F* (*p*(**x**, *t*)), where

**(A.1)** *F*(*p*) ranges in [0, 1] and is a non-negative non-increasing Lipschitz continuous function such that *F* (0) = 1, *F* (1) = 0, *F*(*p*) > 0 for any *p* in (0, 1) and *F*(*p*) = 0 for *p* > 1.

**(A.2)** *K* ≥ 0, 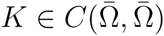 and 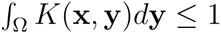.

#### Remarks

1. Notice that system (1.2), (1.3) is not equipped with any boundary conditions. Since it is an integro-differential equation, boundary conditions are implemented by the use of the redistribution kernel *K*. For example, Dirichlet boundary conditions on a bounded domain Ω correspond to kernels that might have support outside of the domain Ω. Then particles moving out of the domain will be lost permanently. Homogeneous no-flux conditions can be employed by requiring that redistribution always happens inside Ω, i.e.

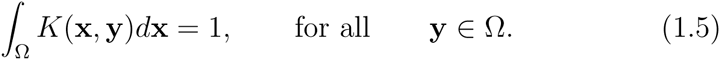
2. Model (1.2), (1.3) falls into the class of birth-jump processes, which were introduced recently in [12]. In a birth-jump process the population growth and population spread are no longer independent processes. On the contrary, it is assumed that newly generated individuals are redistributed instantly. It should be noted that this choice does include standard diffusion terms as special cases. For example if the kernel has the form *K*(| **x** - **y** |) and if *K* is highly concentrated, then a diffusion approximation of the integral can be performed. In fact, this will be done in a forthcoming paper, where we study invasion waves for a reaction-diffusion version of the above model.
3. An advantage of the formulation of the above redistribution kernels is the fact that the quality and local occupancy of the target site can be directly included into the model. If the habitat is inhomogeneous and includes possible uninhabitable patches, then this can be expressed through the choice of *K*(**x, y**). In the context of cancer metastasis, the existence of niches for metastasis is discussed. Niches are favourable environments, where daughter tumors can form [9], and this can be expressed through the kernel *K*(**x, y**).
4. In the original integro-partial differential equation of Hillen et al.[11] the model equations (1.2, 1.3) also contain diffusion terms. If we abbreviate the integral terms as a * product *k* * *u* = *∫*_Ω_ *k*(*x, y, p*(*x, t*))*u*(*y, t*)*dy,* then the corresponding model is

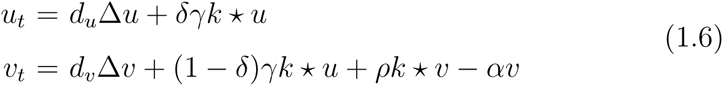

where Ω is a smooth bounded domain. Maddalena [13] studies this model under von Neumann boundary conditions of the form

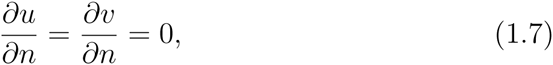

where *n* denotes the outer normal on *∂*Ω. Based on general assumptions on the kernel *k* the integral operators are compact operators on the Banach space *H*^2^(Ω). The system (1.6) appears as heat equation with a compact and Lipschitz continuous perturbation. The solution theory for perturbed parabolic problems is available and Maddalena [13] uses these methods to show results on existence, uniqueness, smoothness and invariant sets. Maddalena did not discuss how the homogeneous von Neumann boundary conditions (1.7) relate to possible boundary flux from the integral terms. It would be an interesting topic for future research to investigate the boundary conditions coming from the integral terms in relation to the classical conditions stemming from the diffusion terms. Since in our model (1.2, 1.3) we have no diffusion terms, we focus on the integral conditions such as (1.5). In our case we study *d*_*u*_ = *d*_*v*_ = 0. Then the leading order terms are the integral terms. The regularity theory for uniform parabolic equations is no longer available and we find it necessary to develop a full solution theory here.

## 2 Existence, uniqueness and boundedness

The above equations (1.2), (1.3) form a non-linear integro-differential equation system. A general existence theory for these type of systems has not yet been developed, hence we present a full solution theory here. We use a fixed-point argument to find unique continuous solutions. First we derive some preliminary estimates, which also show continuous dependence of solutions on the initial conditions and uniqueness. We refine our a-priori estimates to show that solutions stay in the interval [0, 1] for all time, if the initial conditions were in this interval. Finally, in Section 2.2 we construct a contraction mapping whose fixed point is a solution of (1.2), (1.3).

We assume that the initial conditions

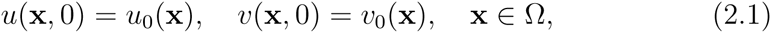

satisfy the following assumptions

**(B.1)** 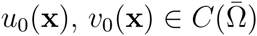,

**(B.2)** 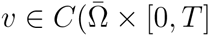.

The existence problem of solutions for (1.2),(1.3) can be stated as:

### Problem (P).

Find a pair *u*(**x**, *t*), *v*(**x**, *t*) such that, for any *T* > 0

- *u, v* ∈ *C*(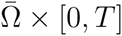);
- *u* ≥ 0, *v* ≥ 0, *u* + *v* ≤ 1, in Ω 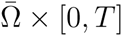;
- *u* and *v* solve the initial value problem (1.2), (1.3), (2.1).

It is quite useful to replace the *v*-equation in (1.3) by an equivalent equation for *p* = *u* + *v*:

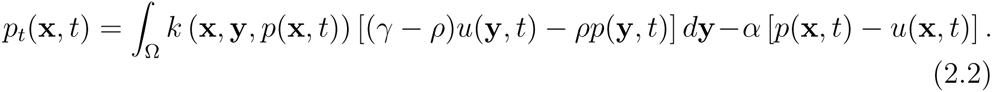

### 2.1 A-priori estimates and uniqueness

We begin with a rough a-priori estimate.

#### Lemma 2.1

*Under assumptions* (A) *and* (B.1) *any solution to (1.2), (1.3), (2.1) is a priori bounded, for bounded t.*

**Proof.** Let us define

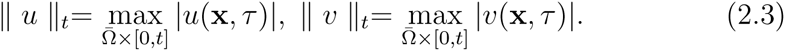

From (1.2), (2.1) we have

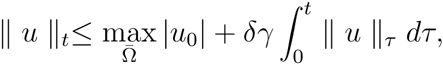

where we used the normalization condition 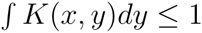. Then from Gronwall’s Lemma we obtain

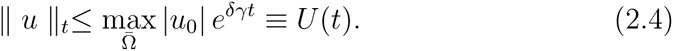

Passing to (1.3), we introduce

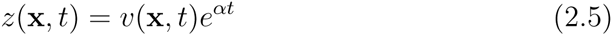

so that

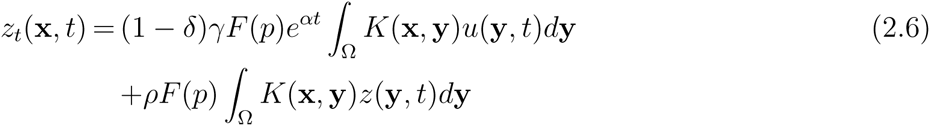

We obtain the estimate

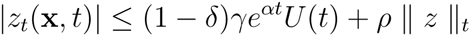

and consequently

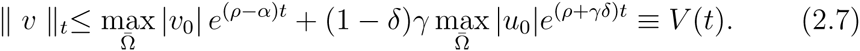

▀

In deriving the above estimates we replaced *F*(*p*) by 1, thus obtaining a rough estimate, which is exponentially growing in time. We will show later that *u* and *v* are indeed uniformly bounded by 1.

At this point we are in a position of proving the following result on continuous dependence on initial conditions:

#### Theorem 2.1

*Let u*^(*i*)^, *v*^(*i*)^ *solve (1.2), (1.3) with data* 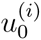, 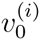, *i* = 1, 2, *and let assumptions* (A) *and* (B.1) *be satisfied. Denote by* Δ*u*, Δ*v*, Δ*p the differences u*^(1)^ — *u*^(2)^, *etc. Then*

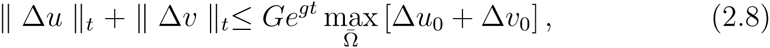

*where G, g are known constants.*

**Proof.** Following the same procedure which we used to get (2.4) we obtain

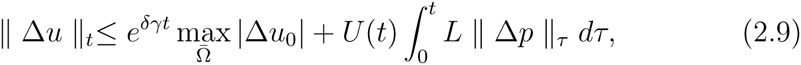

where *L* is the Lipschitz constant of *F*(*p*) and *U* (*t*) (see 2.4) is the larger of the upper estimates for *u*^(1)^ and *u*^(2)^. Moreover, we get

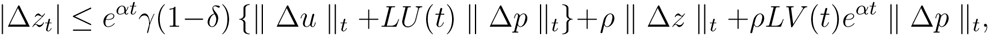

where *z*^(*i*)^ are defined according to (2.5). Thus, ‖ Δ*z* ‖_*t*_ can be estimated via Gronwall’s Lemma and we get

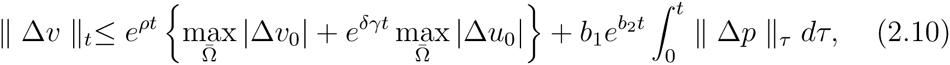

for some known constants *b*_1_, *b*_2_.

Summing (2.9) and (2.10) we get an inequality in terms of Δ*p* and using again Gronwall’s Lemma we conclude the proof.

▀

As a consequence, we have uniqueness:

**Corollary 2.1** *Under assumptions* (A) *and* (B.1) *system (1.2), (1.3) with data (2.1) has at most one solution.*

Sharper estimates on the bounds of *u* and *v* are summarized in the following Lemma:

#### Lemma 2.2

*Assume* (A), (B.1) *are satisfied and let u, v solve equations (1.2), (1.3) in* Ω *×* (*t*_1_, *t*_2_) *for some* 0 ≤ *t*_1_ < *t*_2_.

1. *If u*(**x**, *t*_1_) ≥ 0, *v*(**x**, *t*_1_) ≥ 0 *for all* **x** *∈* Ω, *then*

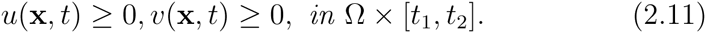
2. *Assume that* 1 *≥ u*(**x**, *t*_1_) *≥* 0, 1 *≥ v*(**x**, *t*_1_) *≥* 0 *for all* **x** *∈* Ω, *then*

i. *if* 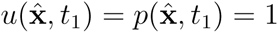, *for some* 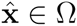, *then* 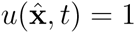, 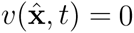, *for t ∈* [*t*_1_, *t*_2_].
ii. *if* 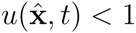, *for some* 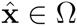, *then* 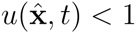, *for t ∈* [*t*_1_, *t*_2_].
3. *If* 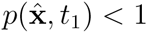, *for some* 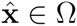, *then* 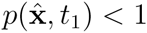, *for t ∈* (*t*_1_, *t*_2_),
4. *If* 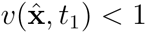, *for some* 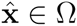, *then* 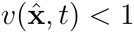, *for t ∈* (*t*_1_, *t*_2_).

**Proof.**

1. Let us start by assuming that

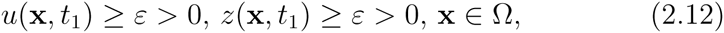

(where *z* is defined by (2.5)), for some *ε* ∈ (0, 1). Let 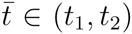 be the first time for which there exists an 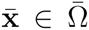 such that 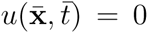. But since (1.2) ensures that *u*_*t*_ ≥ 0 as long as *u* ≥ 0, the solution cannot decay below 0 and *u*(**x**, *t*) *≥* 0. The same argument shows that *z ≥* 0 in Ω × [*t*_1_, *t*_2_], so that *v*(**x**, *t*) ≥ 0. Letting *ε* go to zero and using Theorem concludes the proof of part (1).
2. To show (i) we get from item (1) that *u* ≥ 0. Hence 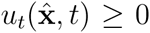 for all *t* ∈ (*t*_1_, *t*_2_) and consequently 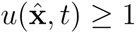, for all *t* ∈ (*t*_1_, *t*_2_). But since item ensures 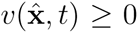 we obtain 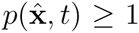 and then 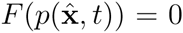 in (*t*_1_, *t*_2_). This implies that *u*_*t*_ = 0 and *u* = 1 in the whole interval. To prove (ii), assume 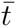 be the first time in (*t*_1_, *t*_2_) such that 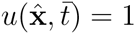. From item (1) we have

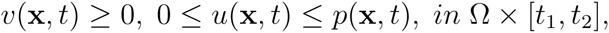

and hence

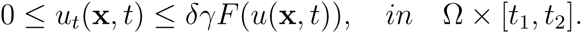 Since *F* is the Lipschitz continuous and *F* (1) = 0, we conclude that the value *u* = 1 cannot be reached in a finite time.
3. We note that the previous results ensure that 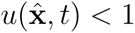, for *t* ∈ (*t*_1_, *t*_2_). Thus, if 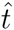 is the first time such that 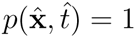, we would have 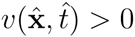 and, because of (2.2), 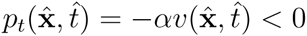, leading to a contradiction.
4. A similar argument as used in item (3) gives item (4).

▀

Now we can apply the above a-priori results to any solution of Problem (P) if (A) and (B) are satisfied, just letting *t*_1_ = 0 in the previous Lemmas. We summarise these facts in the following result on global bounds:

#### Theorem 2.2

*If* (A) *and* (B) *are satisfied, then for any solution of* Problem

(P) *we have*

1. *The “saturated” regions (i.e. regions where p* = 1*) exist only if they exist initially, and never expand.*
2. *Saturated regions shrink if and only if they contain non stem cancers cells.* (3) *All solutions of (1.2), (1.3), (2.1) are solutions of* Problem (P)*. Assumption (B) on the data together with the results of this Section ensure that u ≥* 0, *v ≥* 0, *u* + *v ≤* 1, *in* Ω *×* [0, *T*].

### 2.1 Existence

#### Theorem 2.3

*Under assumptions* (A), (B), Problem (P) *has a unique global solution.*

**Proof.** We use a fixed point argument to show that system (1.2), (1.3), (1.4) has a unique solution (*u, v*) in a subset of the space 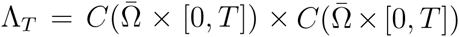 for *T* small enough. Then the results of the previous Section will guarantee that the same pair actually solves Problem (P) (i.e. *u*, *v* are in the physical range) and has all the properties shown there (including continuous dependence on the initial data).

Proceeding as in the previous Section, we already know that, if we prescribe *p*(**x**, *t*) in (1.2), (1.3), the possible solutions of the corresponding system satisfy the a-priori bounds

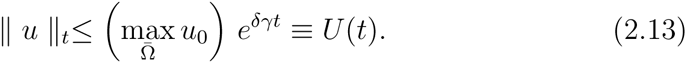

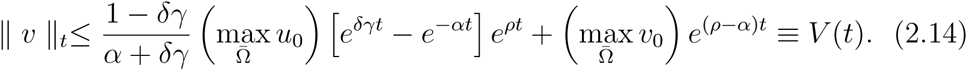

Thus we can take a pair (*u, v*) ∊ Λ_*T*_ such that *u*(**x**, 0) = *u*_0_(**x**), *v*(**x**, 0) = *v*_0_(**x**), and respecting the bounds (2.13), (2.14). We denote such a subset by 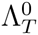. We use *p* = *u* + *v* in (1.2), (1.3) and we look for a solution 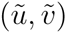 of

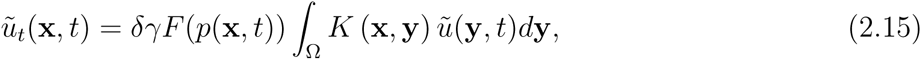

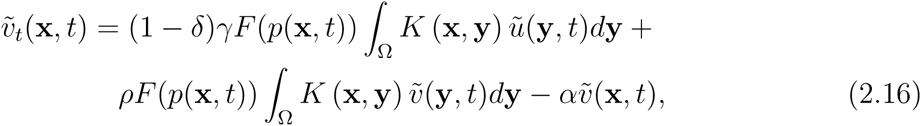

in the same space 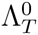.

Since (2.16) is now a linear equation, the existence of the pair 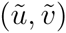 is easily shown using a simple fixed point argument. All we have to do is to analyse the differences 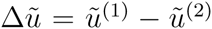, 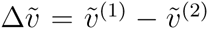 corresponding to the pairs 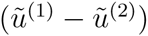, 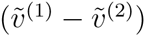 generating the sums *p*^(1)^, *p*^(2)^.

We have that 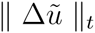, 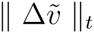 satisfy a system of Gronwall type inequalities with free terms of the form

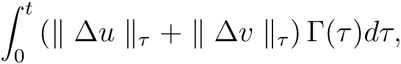

where Γ(*τ*) is either *U* (*τ*) or *V* (*τ*) times some constant. These are the only terms including ‖ Δ*u* ‖ *_t_* and ‖ Δ*v* ‖ *_t_*. Then we can select *T* such that the mapping 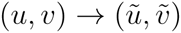 is a contraction in 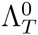.

Once we have proved existence for sufficiently small *T*, we can say that *u*(**x**, *T*), *v*(**x**, *T*) satisfy the same assumptions as *u*_0_(**x**), *v*_0_(**x**). The nonlinearity *F* and the kernel *K* are bounded, therefore the argument can be iterated, yielding global in time existence.

▀

## 3 Numerical examples

We use numerical simulations of the integro-differential equations (1.2), (1.3) to illustrate two effects. First we show that this model shows the tumor-growth paradox. A larger death rate of CC will eventually lead to a larger tumor. This effect was already found in the in silico-model of Enderling [6] and in the ordinary differential equation model of Hillen et al. [11]. Here we confirm that this effect also exists in the integro-differential formulation. We also find that the tumor with larger death rate is stem cell rich, in contrast to the case of a low death rate. We investigate the sensitivity of the tumor growth paradox on the model parameters. We find that this paradox exists for all considered parameter combinations, however in many cases it is not very pronounced.

Secondly, we study one illustrative example of an incomplete radiation treatment. After a rather short time we see that the treated tumor grows larger than it would have been without treatment. Both observations reinforce the importance of the stem cell compartment and its increased resistance to treatment modalities.

Equation (1.2), (1.3) are solved for *x ∊* [- *L, L*], and *t* ∈ [0, 200]. We choose a standard linear volume filling term ([14])

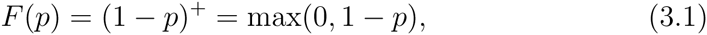

and Gaussian redistribution kernels

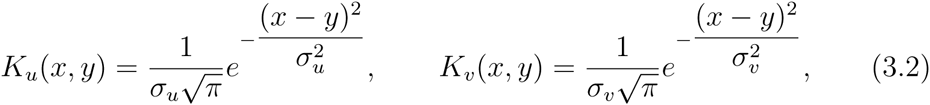

where *σ*_*u*_, *σ*_*v*_ are the standard deviations. To solve system (1.2), (1.3) we used a finite difference scheme, with an explicit forward method in time. The integrals appearing in the equations were computed by the trapezoidal rule.

The initial data were chosen to describe a concentrated tumor mass in the center of the domain which consists of stem cells only:

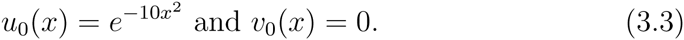

The following parameter values define our standard parameter set

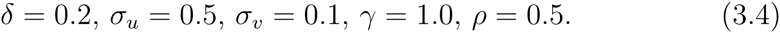

We vary the death rate *α* between two representative values, a small value of *α* = 0.2 and a large value of *α* = 2.

The following examples show the sensitivity of the system to the mortality rate *α* and, in particular, the possible appearance of the tumor growth paradox after a suitable time.

### 3.1 Evidence of the tumor growth paradox

The distribution of *u, v, p* at selected times are reported in Figure 1 for *α* = 0.2 and in Figure 2 for *α* = 2. For small *α* value (*α* = 0.2 in Figure 1) we see that the invasion wave is dominated by CC, whereas in the centre the stem cell population establishes dominance. This behavior is quite different for large *α* = 2 (Figure 2), where the invasion is clearly stem cell dominated and the non-stem cancer cells play only a minor role at the invasion front. To elucidate the appearance of the tumor paradox, we consider the total population of tumor cells, namely

**Figure 1.**
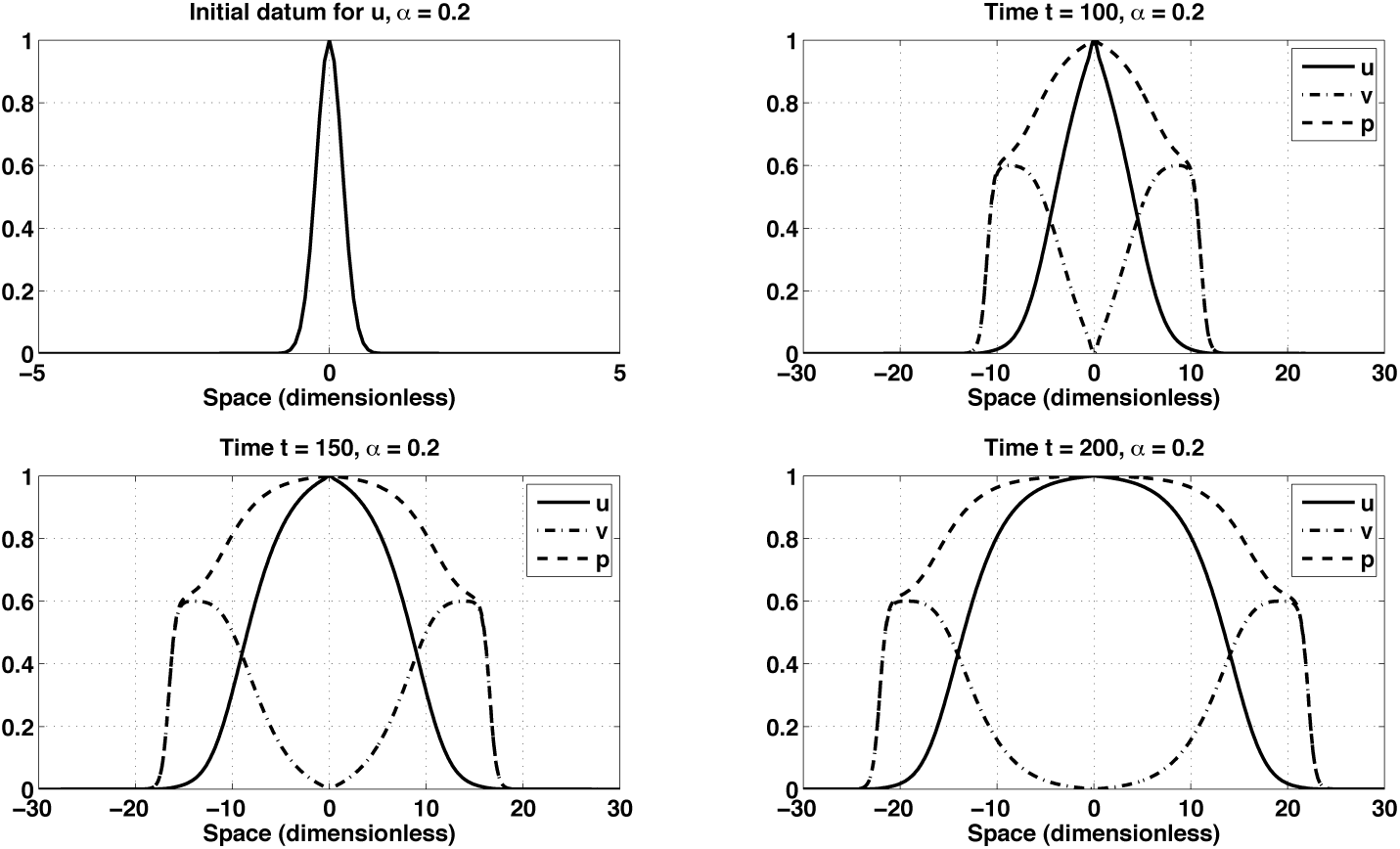
*Plot of u, v, p at selected time instants, with initial conditions (3.3) and parameters from (3.4). Case α* = 0.2*. Detail of space interval* [−30, 30].

**Figure 2.**
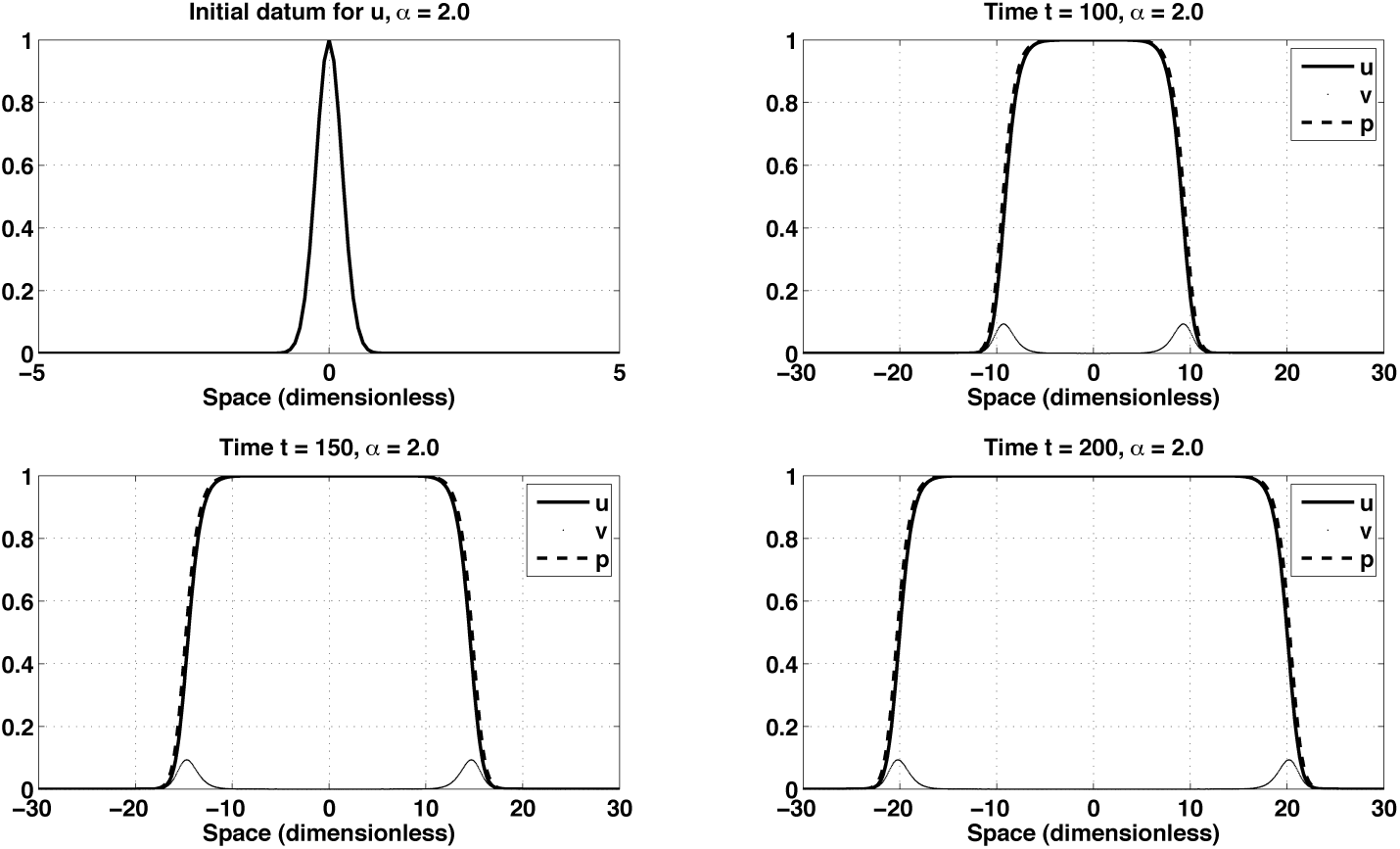
*Plot of u, v, p at selected time instants, with initial conditions (3.3) and parameters from (3.4). Case α* = 2.0*. Detail of space interval* [−30, 30].

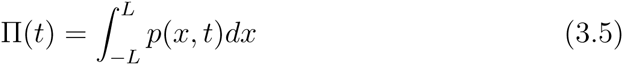

and look for its time evolution, in dependence on *α* (Fig. 3, left). The occurrence of the tumor growth paradox is evident around *t* = 60. The population with a lower death rate *α* initially grows faster, but eventually is exceeded by the populations with the higher *α* value.

**Figure 3.**
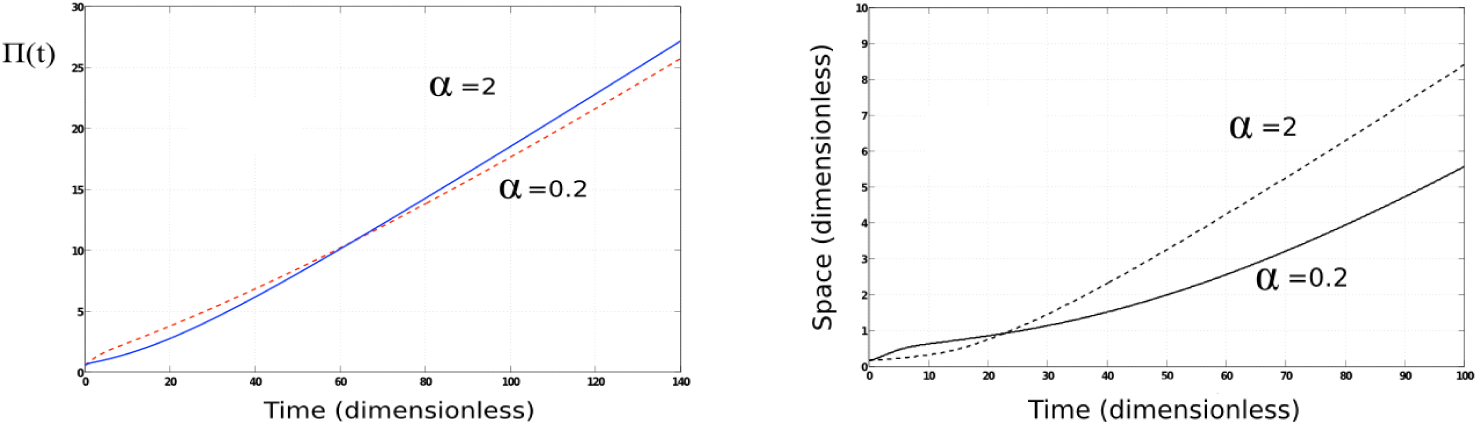
*(Left) Time evolution of total population, Π*(*t*), *for α* = 0.2 *and α* = 2.0*. (Right) Time evolution of x*[*p*=0.8](*t*), *for α* = 0.2 *and α* = 2.0.

To show the tumor growth paradox from a different point of view, the evolution of points tracking a fixed value of *p* was also computed. In Fig.3, right, the 80%- level set of *p* is shown

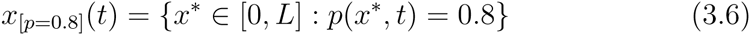

for *α* = 0.2 and *α* = 2. The slope of the corresponding curves is a measure for the invasion speed. Clearly, the curve for larger *α* has a larger invasion speed. We plan to use a forthcoming paper to study the invasion travelling wave speeds in detail.

### 3.2 Dependence on model parameters

To investigate how the tumor growth paradox depends on the model parameters, we investigate a whole range of parameter sets. We found that a faster spread of the stem cells, expressed by an increased the variance *σ*_*u*_, increases the tumour spread, but decreases the tumour growth paradox. By a decreased tumor growth paradox we mean that the difference in population sizes between *α* = 0.2 and *α* = 2 is reduced as compared to models with smaller *σ_u_*. In Figure 4 we compare the simulations at time *t* = 100 for *α* = 0.2 and *σ_u_* = 0.5 (left) and *σ*_*u*_ = 1 (right). The spread in Figure 4 (right) is faster than the spread on the left. Still, the invasion is CC dominated. In the case of *α* = 2 we also observe increased invasion speeds for *σ_u_* = 1, but in this case the invasion is CSC dominated (see Figure 5).

**Figure 4.**
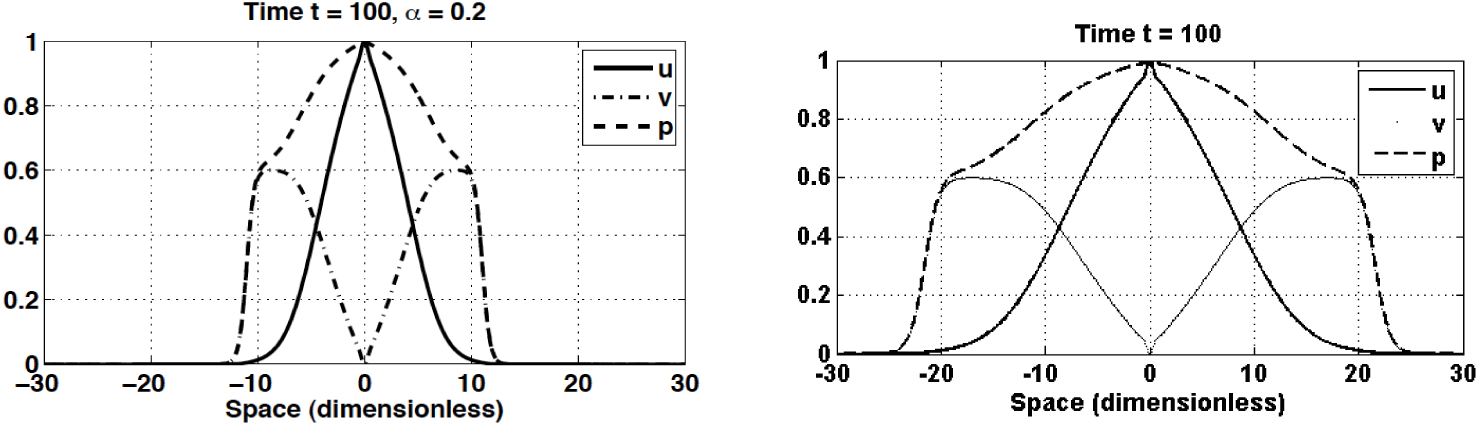
Simulations as in Figure 1 at *t* = 100. Here *α* = 0.2 and *σ_u_* = 0.5 (left) and *σ_u_* = 1 (right). All other parameters are as in (3.4)

**Figure 5.**
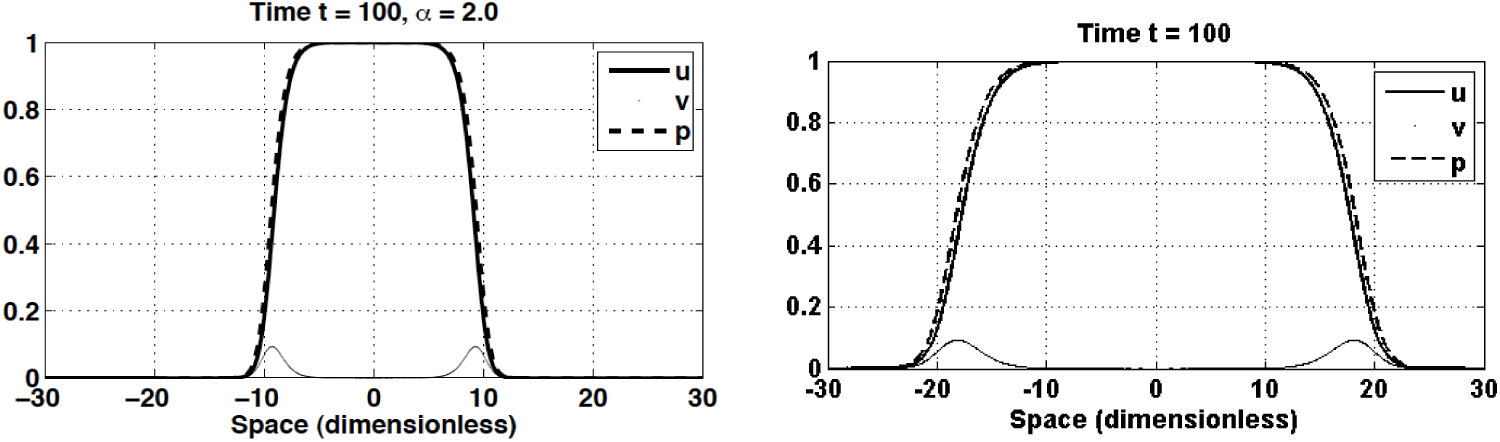
Simulations as in Figure 1 at *t* = 100. Here *α* = 2 and *σ_u_* = 0.5 (left) and *σ_u_* = 1 (right). All other parameters are as in (3.4)

A variation of the spread rate for *v, σ_v_*, only has a minimal effect on the simulation results (not shown). However, a variation of the rate of CSC renewal *δ* makes a big difference. In Figure 6 we increase *δ* from 0.2 to 0.5. We show the simulation for *t* = 100 and *α* = 0.2 on the left and *α* = 2 on the right. In both cases the invasion is driven by the CSC compartment. The tumor growth paradox still arises, as seen in Figure 7 (right), but the difference in the total populations is negligible for *δ* = 0.5.

**Figure 6.**
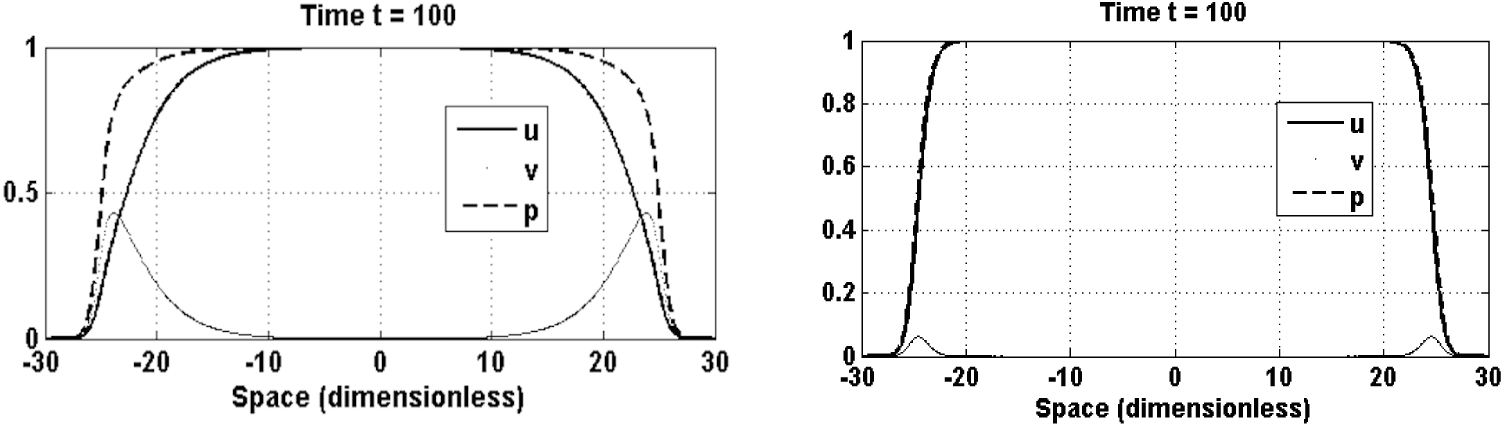
Simulations as in Figure 1 at *t* = 100. Here *δ* = 0.5 and *α* = 0.2 (left) and *α* = 2 (right). All other parameters are as in (3.4)

**Figure 7.**
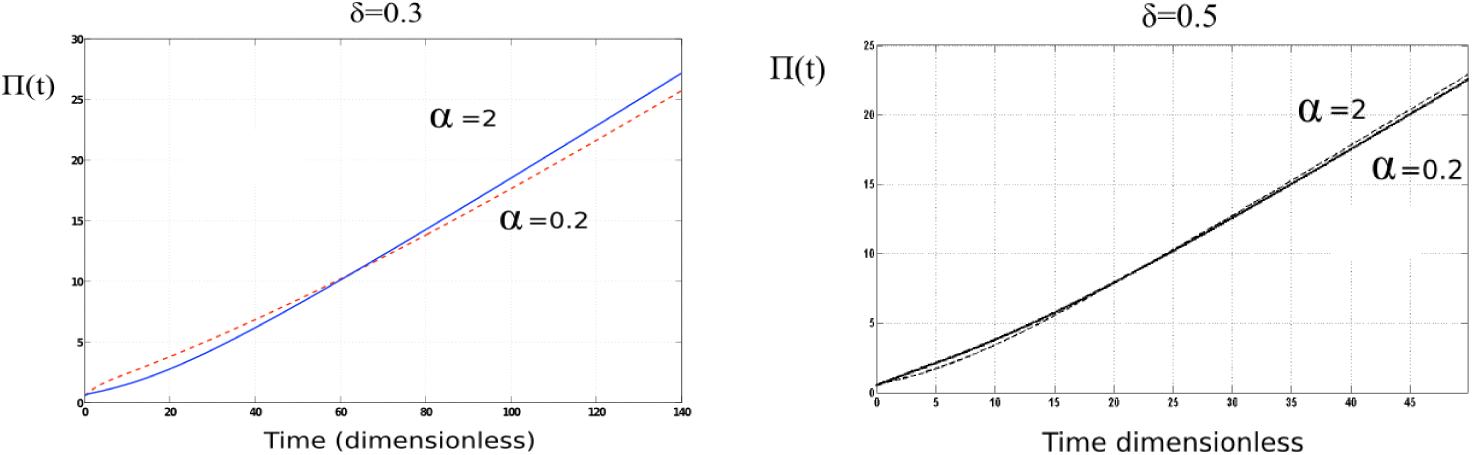
Time evolution of the total population π(*t*) for *δ* = 0.3 (left) and *δ* = 0.5 (right). All other parameters are as in (3.4)

### 3.3 Simulating the effect of a radiation treatment

To illustrate the basic effect of stem cell resistance to radiation, we use a rather ad-hoc approach. We consider one radiation dose (at time *t*\* = 50 in this example), and we denote the reduction of the cell population by a factor 1 *ϕ, ϕ* [0, 1]. We are aware of detailed radiation treatment models involving the linear quadratic model and tumor control probabilities, but a detailed discussion of this framework does not add to the argument we want to make here. For details on radiation treatment modelling we refer to [8] and [3]. Here, at the treatment time *t\**, we choose new initial data as

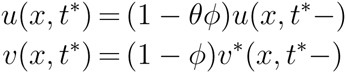

where 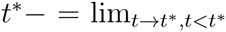. The term (1 − *ϕ*) denotes the surviving fraction of CC while (1 − *θϕ*) denotes the larger surviving fraction of CSC. The value of *θ*, 0 *≤ θ ≤* 1, describes radio resistance of stem cells.

Results reported in Fig. 8 correspond to *ϕ* = 0.95, *θ* = 0, for *α* = 0.2, as selected times after treatment. Other model parameters have value as in (3.4). The upper figure shows the effect of radiation at *t* = 50. The solid line shows the tumor distribution *p*(*x, t*) just before treatment, and the dashed line just after treatment. The reduction is lowest in the centre, where most CSC are residing. The lower figure in Figure 8 shows the tumor evolution at time *t* = 150, where the solid line (without treatment) basically coincides with the dashed line (with treatment). Hence incomplete treatment seems to have no effect on the tumor.

**Figure 8.**
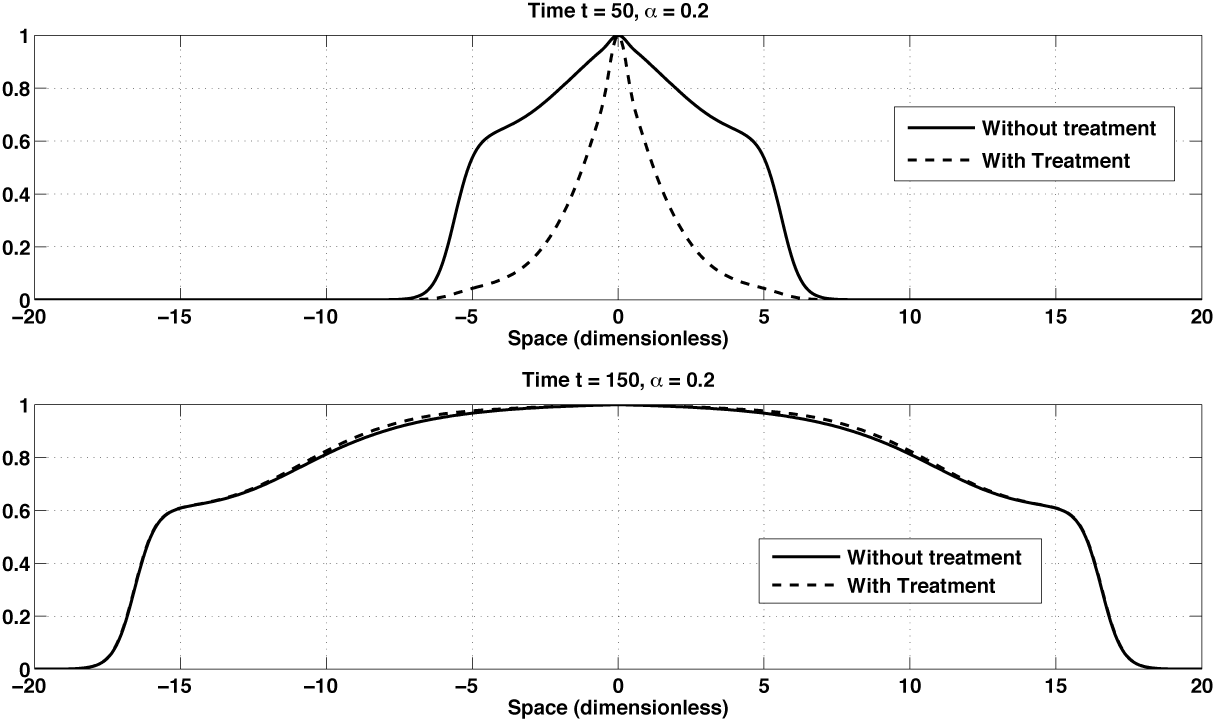
*Plot of p at selected time instants in case of radiation treatment at t* = 50*. Case α* = 0.2*. Detail of space interval* [−20, 20].

In Fig. 9 we show the total population on the left and the level set *p* = 0.8 on the right. Here we see that the progression with treatment (dashed line) is slightly faster than without treatment (solid line). Again, a mild tumor growth paradox is at play.

**Figure 9.**
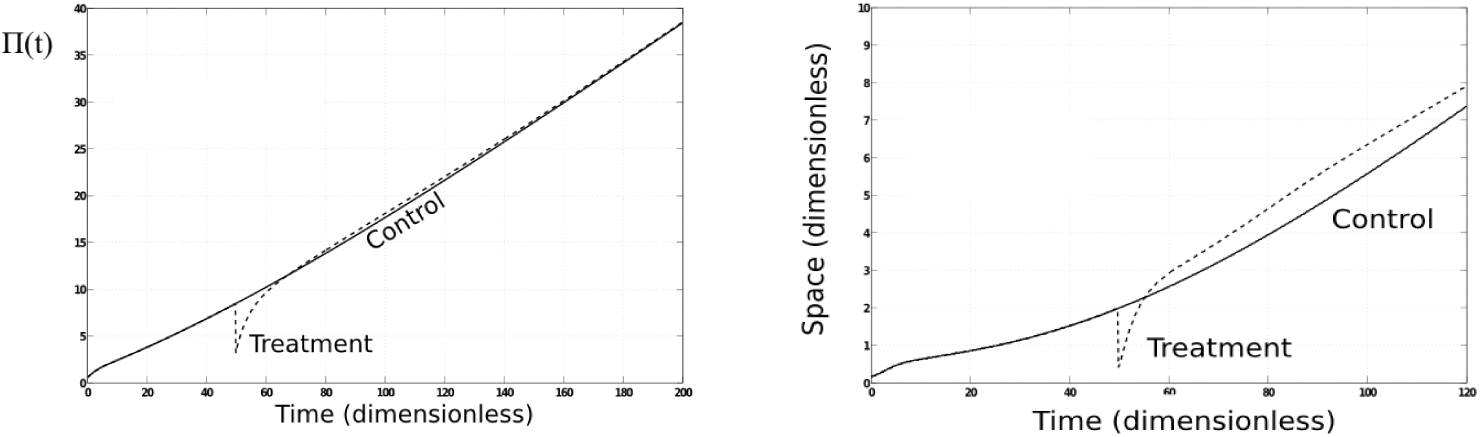
*(Left) Time evolution of total population, Π* (*t*), *for ϕ* = 0.95 *and α* = 0.2, *(Right) Time evolution of x*[*p*=0.8](*t*), *for ϕ* = 0.95 *and α* = 0.2.

## 4 Discussion

The increased resistance of cancer stem cells to various treatment modalities, as compared to non stem CC, is one of the major obstacles of cancer treatment. Cancer stem cells have been used to explain many observed phenomena of cancer progression, including the tumor growth paradox. Our model extends previous results of the tumor growth paradox. The individual based model of Enderling [6], the ODE of Hillen [11] and our system (1.2), (1.3) make it perfectly clear, that, as soon as CSC an CC compete for space and resources, a tumor growth paradox exists.

We have studied an ad hoc radiation treatment with a single application of radiation (one fraction) and we found that incomplete treatment leads to a selection of CSC. We could include more realistic treatments, or chemotherapy and other treatments, but the principle will be the same. Any stress applied to the tumor will lead to a selection of CSC, and this is the point we wanted to make here. The consideration of more realistic treatments is left to further studies. It is clear, however, that a successful treatment needs to target CSC. One such strategy was suggested by Youssefpour, Lowengrub et al. [18]. They suggest that conventional treatment, such as radiation treatment, should be combined with a differentiation therapy. Differentiation therapy describes a chemotherapy, where the chemotherapeutic agent pushes stem cells into the differentiation cascade. Several candidate drugs are known and currently tested. In [1] the above ODE stem cell model (1.1) was used to test Lowengrub’s hypothesis for specific cancers (secondary brain tumors, breast cancer, and head and neck cancer). It was confirmed that for brain and head and neck cancers a combination therapy can reduce the necessary radiation dose, while the benefit for breast cancer treatment was limited (according to the mathematical model).

As outlined before, our cancer stem cell model (1.2), (1.3) falls into the class of birth-jump models as introduced in [12]. The existence theory developed here is only a start. Many more interesting mathematical questions lie ahead for future research, including models for invasion, travelling waves and spatial pattern formation.

## Acknowledgements

We are grateful to detailed comments from two anonymous referees. TH is supported through NSERC.

## References

[1] J. Bachman and T. Hillen. Mathematical optimization of the combination of radiation and differentiation therapies of cancer. Frontiers in Molecular and Cellular Oncology, 2012. free online, doi: 10.3389/fonc.2013.00052.

[2] E. Beretta, V. Capasso, and N. Morozova. Mathematical modelling of cancer stem cells population behavior. Math. Modelling of Natural Phenomena, 7(1):279–305, 2012.

[3] A. Dawson and T. Hillen. Derivation of the tumour control probability (TCP) for a cell cycle model. Comput. and Math. Meth. in Medicine, 7:121–142, 2006.

[4] D. Dingli and F. Michor. Successful therapy must eradicate cancer stem cells. Stem Cells, 24(12):2603–2610, 2006.

[5] T. Dittmar and K.S. Zänker. Role of Cancer Stem Cells in Cancer Biology and Therapy. CRC Press, Boca Raton, FL, USA, 2013.

[6] H. Enderling, A.R.A. Anderson, Chaplain M.A.J., A. Beheshti, L. Hlatky, and P. Hahnfeldt. Paradoxical dependencies of tumor dormancy and progression on basic cell kinetics. Cancer Research, 69(22):8814–8821, 2009.

[7] R. Ganguli and I.K. Puri. Mathematical model for the cancer stem cell hypothesis. Cell Proliferation, 39:3–14, 2006.

[8] J. Gong, M. M. dos Santos, C. Finlay, and T. Hillen. Are more complicated tumor control probability models better? Math. Med. Biol., 30(1):1–19, 2011.

[9] D. Hanahan and R.A. Weinberg. Hallmarks of cancer: The next generation. Cell, 144:646–674, 2011.

[10] G. Hek. Geometric singular perturbation theory in biological practice. J. Math. Biology, 60(3):347–386, 2010.

[11] T. Hillen, H. Enderling, and P. Hahnfeldt. The tumor growth paradox and immune system-mediated selection for cancer stem cells. Bull. Math. Biology, 75(1):161–184, 2013.

[12] T. Hillen, B. Greese, J. Martin, and G. de Vries. Birth-jump processes. J. Biol. Dynamics, 2014. doi: 10.1080/17513758.2014.950184.

[13] L. Maddalena. Analysis of an integro-differential system modelling timor growth. Appl. Math. Comp., 245:152–157, 2014.

[14] K.J. Painter and T. Hillen. Volume-filling and quorum-sensing in models for chemosensitive movement. Canadian Appl. Math. Quart., 10(4):501–543, 2002.

[15] R.V. Sole, C. Rodriguez-Caso, T.S. Diesboeck, and J. Saldance. Cancer stem cells as the engine of unstable tumor progression. J. Theor. Biol., 253:629–637, 2008.

[16] T. Stiehl and A. Marciniak-Czochra. Mathematical modeling of leukemogenesis and cancer stem cell dynamics. Math. Modelling of Natural Phenomena, 7:166–202, 2012.

[17] Z. Sun and N.L. Komarova. Stochastic control of proliferation and differentiation in stem cell dynamics. Math. Biosci., 240:231–240, 2012.

[18] H. Youssefpour, X. Li, A.D. Lander, and J.S. Lowengrub. Multispecies model of cell lineages and feedback control in solid tumors. Journal of Theoretical Biology, 304(0):39–59, 2012.

